# The cell cycle regulates cell-penetrating peptide cytosolic uptake via differential expression of KCNH1 potassium channel

**DOI:** 10.64898/2026.09.01.748619

**Authors:** Francisco Tomás Ribeiro, Ali Hallaj, Roger Mora, Anita Lüthi, Evgeniya Trofimenko, Christian Widmann

**Affiliations:** Department of Biomedical Sciences, University of Lausanne, 1005 Lausanne, Switzerland; Department of Fundamental Neurosciences, University of Lausanne, 1005 Lausanne, Switzerland

**Keywords:** Cell-penetrating peptides, direct translocation, cell cycle, membrane potential, fluorophores, D-amino acid, KCNH1

## Abstract

Membrane potential fluctuations, driven by ion channel activity, are important regulators of processes such as action potential propagation in neurons and muscle cell contraction. These variations also occur in non-excitable cells, influencing various cellular processes, including, but not limited to, direct translocation of cell-penetrating peptides (CPPs) into the cytosol. CPPs are cationic peptide vectors that facilitate the delivery of cargos into cells. The efficiency of CPP direct translocation varies significantly within the same cell population, a phenomenon that is not understood mechanistically.

In this study, we used an unbiased transcriptomic approach to identify genes involved in efficient CPP cytosolic uptake. This revealed a close correlation between the expression of cell cycle-related genes and the ability of cells to take up CPPs by direct translocation. Monitoring phases of the cell cycle with the PIP-FUCCI system showed that CPP direct translocation was most efficient in the G2/M and S phases and least efficient during the G1 phase. This was correlated with the cells’ plasma membrane potential: cells in G2/M and S phases being more polarized than cells in the G1 phase. The expression of the KCNH1 voltage-gated potassium channel was found to be highest in the phases of the cell cycle most permissive for CPP direct translocation. Pharmacological inhibition of KCNH potassium channels efficiently blocked CPP direct translocation in these cell cycle phases. These data indicate that KCNH1 controls the cell membrane potential in a cell cycle-dependent manner and consequently the ability of cells to take up CPPs by direct translocation. Our findings highlight the importance of the cell cycle in regulating CPP uptake. Variations in the plasma membrane potential along the cell cycle should be considered in the development of CPP-based therapeutics and their applications.

## Introduction

Changes in membrane potential, brought about by the activity of ion channels, are the drivers of action potential generation and transmission in neurons, and contraction in muscle cells [1]. Membrane potential variations also occur in non-excitable cells and can impact several cellular processes, such as the cytosolic uptake of cell penetrating peptides (CPPs) by direct translocation [2–6].

CPPs are short, cationic molecules of 5 to 30 amino acids that can enter cells via two non-mutually exclusive mechanisms: direct translocation where CPPs can directly access a cell’s cytosol by crossing the cell membrane; or through endocytosis, which requires the formation of an endocytic vesicle to bring CPPs into cells [5, 7–10]. Most used CPPs in research are TAT (Trans-Activator of Transcription) from HIV-1 [7, 9–11], Penetratin, originating from the Antennapedia homeodomain [7, 9, 10, 12], and synthetic arginine- and tryptophane-rich molecules [13, 14]. CPPs can be used as delivery vectors to bring cargos into cells [7, 10]. Several CPP-conjugated compounds are currently being evaluated in phase II and III clinical trials [15–17].

We and others have provided evidence that CPP direct translocation into cells occurs through water pores [5, 18–22]. Initially, cationic peptides interact with the negatively charged components of the cell membrane, such as glycosaminoglycans and phospholipids [9]. These interactions allow CPPs to accumulate at the cell membrane. The cationic nature of CPPs, in combination with the activity of potassium channels, lead to the accumulation of positive charges at the surface of cells. This locally hyperpolarizes the membrane, in turn, increasing the probability of water pore formation. The water pores are then used by the CPPs to access the cell’s cytosol [5]. Once the CPPs cross the plasma membrane, the positive charges carried by them dissipate the electrochemical gradient, which causes the water pores to collapse. Due to the transient existence of water pores, CPP translocation across the plasma membrane does not, or only minimally, affect cell viability. Naturally hyperpolarized cells such as neurons, require lower CPP concentrations for direct translocation to be induced compared to, for example, immortalized cells in culture [5].

A phenomenon that is commonly observed by researchers working on CPPs is that cells within a given population can have strikingly different abilities to take up CPPs, ranging from no uptake to saturation of the cytosolic signal [3, 5, 23]. Although various mechanisms have been proposed to describe the initial CPP uptake into cells, there is currently no explanation for the variations in the efficiency of CPP internalization in a cell culture.

Here, we show that the ability of cells to take up CPPs varies along the cell cycle because of differential expression of the KCNH1 potassium channel and the ensuing modulation of the plasma membrane potential.

## Results

### Correlation between expression of cell cycle-related genes and extent of CPP uptake by direct translocation

CPP uptake efficiency may depend on various factors including, but not limited to, cell membrane composition, CPP sequence and CPP concentrations. *Figure 1 – Figure Supplement 1* shows how CPP concentration affects the mode of entry and how this is dependent on the plasma membrane potential. Strikingly, beyond these factors, within the same population of cells, we and others have observed a significant heterogeneity in CPP cytosolic internalization, ranging from non-uptakers to cells with saturated CPP cytosolic signal [3, 5, 23]. This is for example seen in U2OS cells incubated with 15 µM of the Cy5-tagged nona-arginine CPP (Cy5-R9) (*Figure 1A*). To identify genes playing a role in modulating CPP entry into cells, we performed a transcriptomic analysis on cells with differential CPP uptake by sorting the 5% dimmest and 5% brightest cells, corresponding to cells with no/low direct translocation and efficient direct translocation, respectively (*Figure 1 – Figure Supplement 2A*). Gene Set Enrichment Analysis (GSEA) [24, 25] of the transcriptomics data was used to establish which specific sets of genes were altered between the two analysed conditions. Most gene sets enriched in cells with high CPP uptake were related to mitosis and DNA replication, as well as other aspects of cell cycle regulation (*Figure 1B* and *Figure 1 – Figure Supplement 2B*). For example, the mRNA encoding the CDK1 cyclin-dependent kinase was enriched in the cell subpopulation containing high levels of cytosolic CPPs (*Figure 1C and Figure 1 – Figure Supplement 2C*). Other examples include mRNAs encoding cyclin F and cyclin A2, master regulators that drive cells through critical stages of cell division (*Figure 1C and Figure 1 – Figure Supplement 2C*). These data indicate that the cell cycle progression plays a role in modulating the heterogeneity of CPP cytosolic uptake within a cell population.

**Figure 1.**
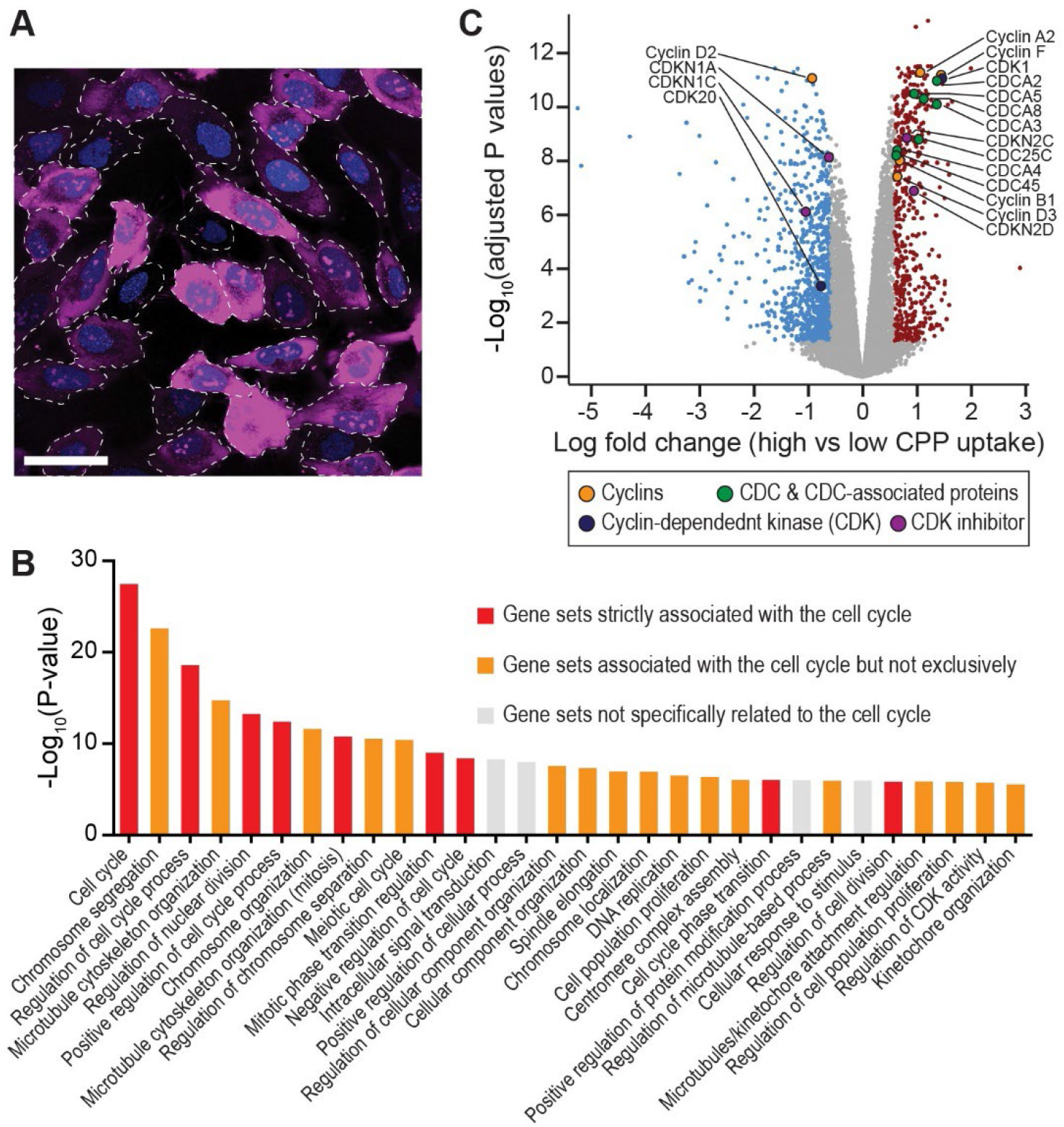
Cell cycle-related genes are differentially expressed in cells with high vs low CPP cytosolic uptake. **(A)** CPP cytosolic uptake is heterogenous within a cell culture. Representative confocal image of live U2OS cells incubated for 5 minutes at 37°C with 15 µM Cy5-R9, followed by PBS washes. The dotted lines represent the cell contours. Scale bar: 50 µm. **(B)** Gene ontology enrichment analysis performed on the transcriptomics data of U2OS subpopulations with high vs low Cy5-R9 uptake. **(C)** Volcano plot (fold-change cut-off = 1.5; p-value cut-off = 0.05) generated from the transcriptomics data of cells with high vs low Cy5-R9 cytosolic uptake. Examples of differentially expressed regulators of the cell cycle are highlighted.

### CPP direct translocation is least efficient during the G1 and M phases

To study the relationship between the cell cycle and CPP uptake, we took advantage of the U2OS PIP-FUCCI system [26] that allows the monitoring of cell cycle progression via three different fluorophore-tagged cell cycle-regulated markers that can be detected by microscopy and flow cytometry (*Figure 2 – Figure Supplement 1 A-B*). Of note, no bleed-through from the CPP signal was detected in the channels used to detect the three markers of the PIP-FUCCI system (*Figure 2 – Figure Supplement 2*). As the PIP-FUCCI system cannot discriminate the G2 from the M phase, nocodazole, a microtubule-disrupting agent [27], can be used to block cells at the onset of mitosis and hence synchronize cells in early mitosis (*Figure 2 – Figure Supplement 3A*). Although CPP uptake could be detected in all cell cycle stages, its efficiency was lowest in G1 and absent in M (*Figure 2* and *Figure 2 – Figure Supplement 3B-C*). On the other hand, CPP direct translocation was favoured in S and G2 (*Figure 2*). This is in line with our transcriptomics data showing that cells with the capacity for highest CPP cytosolic uptake were enriched in genes expressed during the transition to S phase (*Figure 1B-C and Figure 1 – Supplement 1B-C*). Our data indicate that cells in different phases of the cell cycle have distinct abilities to take up CPPs by direct translocation. On the other hand, CPP uptake by endocytosis was not affected throughout the different cell cycle stages (*Figure 2 – Figure Supplement 4A*).

**Figure 2.**
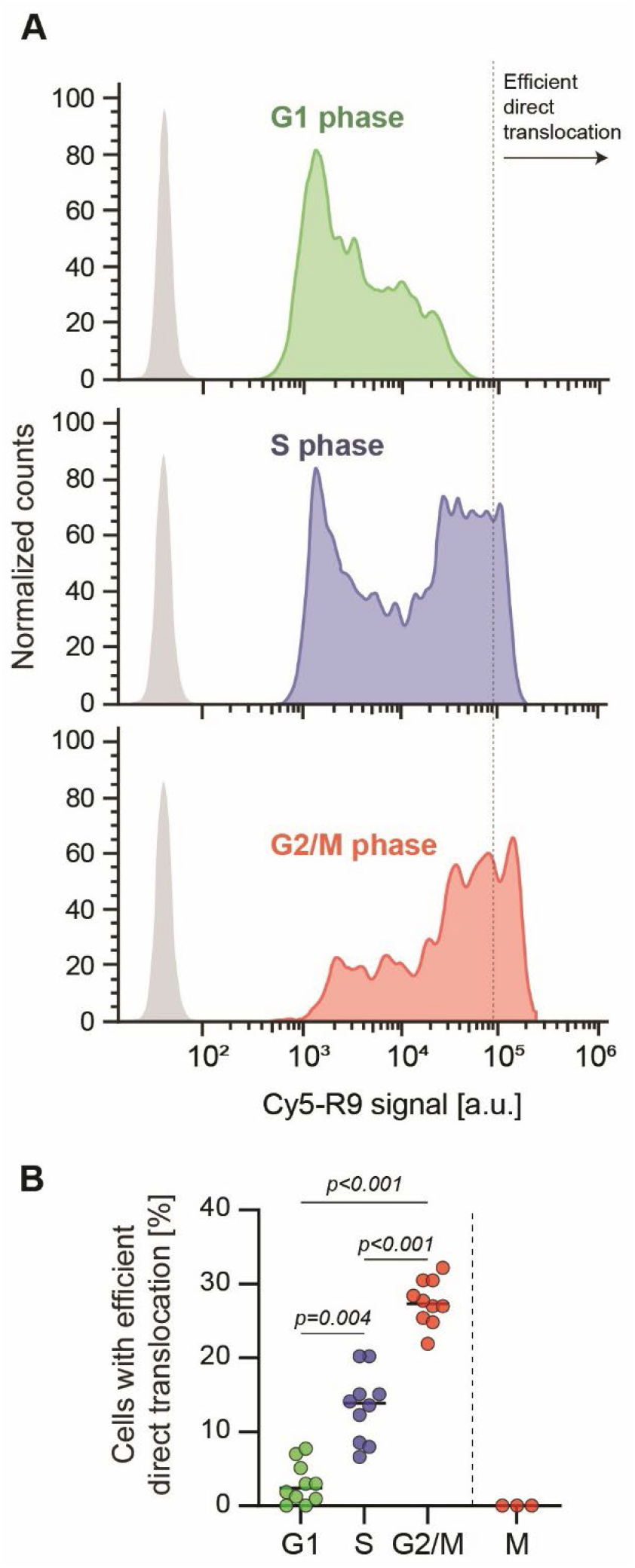
CPP direct translocation is least efficient during the G1 phase of the cell cycle. **(A)** CPP direct translocation varies across the cell cycle. Representative flow cytometry profiles of Cy5-R9 uptake in PIP-FUCCI U2OS cells in the indicated phases of the cell cycle. Cells were incubated with 15 μM Cy5-R9 for 5 minutes at 37°C. Cell counts were normalized. A total of 10 000 cells were analysed in each condition (n=10). The grey profiles correspond to control cells not incubated with Cy5-R9. **(B)** Percentage of cells within a given cell cycle stage that acquired Cy5-R9 through efficient direct translocation as defined in panel A. Efficient direct translocation in M phase cells (data on the right of the by dashed line) was quantitated from confocal microscopy data displayed in *Figure 2 – Figure Supplement 3* (n=3).

### Membrane potential changes across the cell cycle correlate with the extent of CPP direct translocation

Direct translocation of CPPs into cells requires plasma membrane hyperpolarization[2–6, 18–21, 28]. When cells are depolarized, CPP translocation across the cell membrane is inhibited [2, 3, 5] but CPP endocytosis proceeds normally [29] (*Figure 1 – Figure Supplement 1 and Figure 2 – Supplement 4A*). Therefore, we hypothesize that cells with least efficient CPP direct translocation, i.e. cells in the G1 and M phase, are more depolarized than cells in S phase or G2/M transition phase. We measured the resting membrane potential of cells across the cell cycle stages by whole-cell patch clamp (*Figure 3A*). These experiments were performed on double thymidine-synchronized PIP-FUCCI cells that were then released for different periods of time to position cells in given cell cycle stages. The identity of the cell cycle stages was confirmed by flow cytometry using the PIP-FUCCI markers or via DNA content measured by Hoechst staining (*Figure 2 – Figure Supplement 4C-D)*.

**Figure 3.**
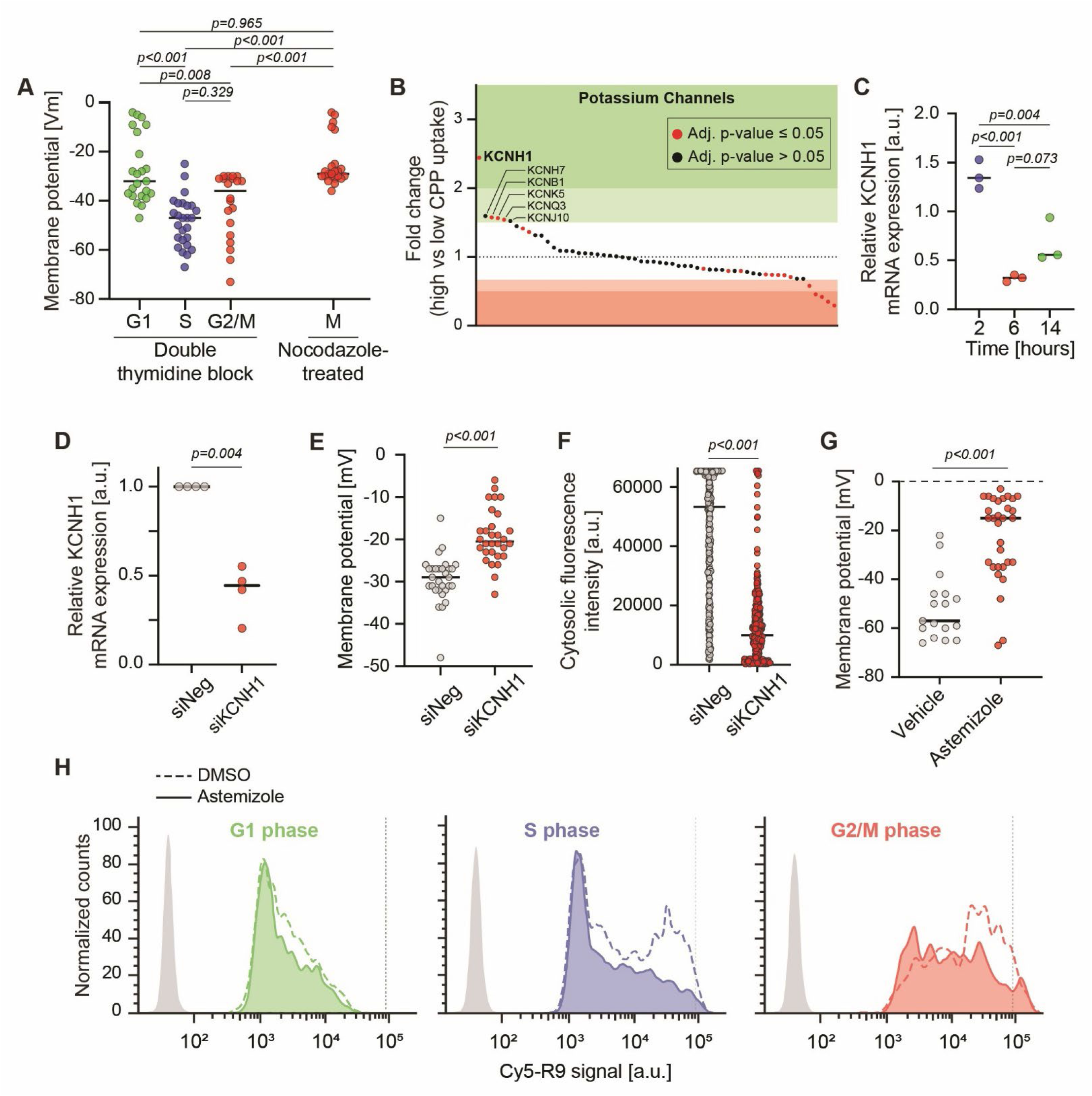
The KCNH1 potassium channel controls CPP direct translocation. **(A)** Membrane potential varies across the cell cycle. Membrane potential measurements were performed on thymidine-synchronized cells at different time points corresponding to G1, S, and G2/M phases; and in nocodazole-arrested cells (M phase). Resting membrane potential measurements were acquired by whole-cell patch clamp. **(B)** Differential expression of potassium channels in cells with high vs low CPP cytosolic uptake. Potassium channels were ranked along the x axis according to the fold change of their mRNA expression in cells with high vs low CPP uptake. **(C)** KCNH1 mRNA levels vary across the cell cycle. Semi-quantitative PCR assessment of KCNH1 mRNA levels in double thymidine-synchronized U2OS cells at different timepoints post release. Samples were normalized to GAPDH mRNA levels and to the 0 hour post double thymidine block release. Statistical analysis was performed using ANOVA multiple comparison. **(D)** Semi-quantitative PCR assessment of KCNH1 mRNA levels in control (siNeg) and KCNH1 knocked down (siKCNH1) U2OS cells. Samples were normalized to GAPDH mRNA levels and to the control values (n=4). Statistical analysis was performed using paired t-test. **(E)** KCNH1 depletion leads to resting cell membrane potential depolarization. Membrane potential was assessed with whole-cell patch clamp in control (siNeg) and KCNH1-knocked down (siKCNH1) in U2OS cells (n=3). Statistical analysis was performed using paired t-test. **(F)** KCNH1 knock-down reduces R9-Cy5 direct translocation. A total of 326 cells were analysed in three independent experiments. Statistical significance was performed using Welch’s t test. **(G)** U2OS cells are more depolarized when KCNH1 is acutely inhibited. Cells were preincubated with 10 µM astemizole or with vehicle (0.1% DMSO) for 5 minutes at 37°C and membrane potential was then measured. Statistical analysis was performed using Welch’s t-test. **(H)** Astemizole prevents efficient CPP direct translocation in cells at the S and G2/M stages. PIP-FUCCI cells were incubated for 5 minutes at 37°C with 10 µM astemizole or vehicle (0.1% DMSO) in the presence of 15 µM Cy5-R9. Cells were then analysed by flow cytometry. The value where the dashed line crosses the x axis was used for defining the quantitation shown in *Figure 3 Supplement 2*.

The cell membrane potential measured in cells in G2/M varied from -75 to -30 mV, with a distinct cluster at around -35 mV (Figure 3A). Our interpretation is that cells at the G2 onset are hyperpolarized to values as low as -75 mV. Then, as they progress toward mitosis, they get depolarized, eventually reaching values around -35 mV. The resting membrane potential of mitotic cells, synchronized with nocodazole treatment, overlapped with the membrane potential of the most depolarized cells in G2 (*Figure 3A*). As expected from this depolarized state, cells in early M phase were inefficient at taking up CPP by direct translocation (*Figure 2* and *Figure 2 – Supplement 3*).

Cells in G1 were found to be more depolarized than cells in S or G2/M transition, with the depolarized values overlapping with those in M phase (*Figure 3A*). Some G1 cells indeed had strongly depolarized membrane potentials (-10 to -5 mV). We interpret these values as corresponding to post-mitotic cells that just entered G1 (*Figure 3A)*. As the cells start the transition from G1 to S phase, they become progressively hyperpolarized, which aligns with the CPP uptake data, showing that direct translocation in favoured in S phase (*Figure 2*).

Taken together, these data indicate that the membrane potential varies along the cell cycle which in turn impacts the ability of cells to acquire CPP by direct translocation (see the model displayed in Figure 3B).

### The cell cycle-dependent KCNH1 potassium channel promotes efficient CPP direct translocation

Membrane potential is set by the electrochemical gradient of potassium, sodium, chloride and calcium ions. However, due to the high permeability of potassium ions, potassium channels play a major role in the regulation of the cell membrane potential and, indirectly, the ability of cells to take up CPPs by direct translocation [2–5, 18–22]. Therefore, we sought to determine whether the measured changes in membrane potential across the cell cycle (*Figure 3A*), could be accompanied by cell cycle-dependent changes in expression of specific potassium channels. Based on our transcriptomics performed in U2OS cells, KCNH1 (Kv10.1), a voltage-activated potassium channel, stood out, with its mRNA being enriched in the high CPP cytosolic uptake subpopulation (*Figure 3B*) and in the cell cycle stage most permissive for CPP direct translocation, i.e. the S phase (*Figure 3C*). In LAN1 cells, expression of KCNH1 also correlated with the ability of cells within a given cell stage to take up Cy5-R9 by direct translocation (*Figure 3 – Supplement 1A and C*). The same was not observed in HEK293T cells, where KCNH1 expression does not depend on cell cycle progression (*Figure 3 – Supplement 1B and D*).

To directly determine if KCNH1 modulates the plasma membrane potential, we reduced its expression using an siRNA-based approach (*Figure 3D*). This resulted in a 10-mV membrane depolarization (*Figure 3E*), which in turn led to a significant decrease in CPP direct translocation visualized by confocal microscopy (*Figure 3F)*. Similar results were obtained using TAT instead of R9 CPP (*Figure 3 – Supplement 2*). Acute pharmacological inhibition of KCNH1, using astemizole, [1, 30–32], also triggered cell membrane depolarization and inhibition of Cy5-R9 direct translocation in S and G2/M phases (*Figure 3 – Figure Supplement 2*).

In conclusion, KCNH1, a voltage-gated potassium channel with differential expression across the cell cycle, impacts cell membrane potential. This, in turn, affects CPP internalization via direct translocation in a cell cycle dependent manner.

## Discussion

In the present work, we show that CPP direct translocation efficiency varies along the cell cycle in a KCNH1-dependent manner. Our data support the following model (*Figure 4A*). During the G1 stage, cells lower their membrane potential (hyperpolarization) progressively and continue to do so into S phase, reaching a minimum in late S or at the S-G2 border. During G2/M progression, the membrane potential increases steadily until strong depolarization is achieved at the onset of mitosis [33], corroborated by our data of cells synchronized in early mitosis. Membrane potential oscillations are inversely proportional to CPP direct translocation across the cell membrane.

**Figure 4.**
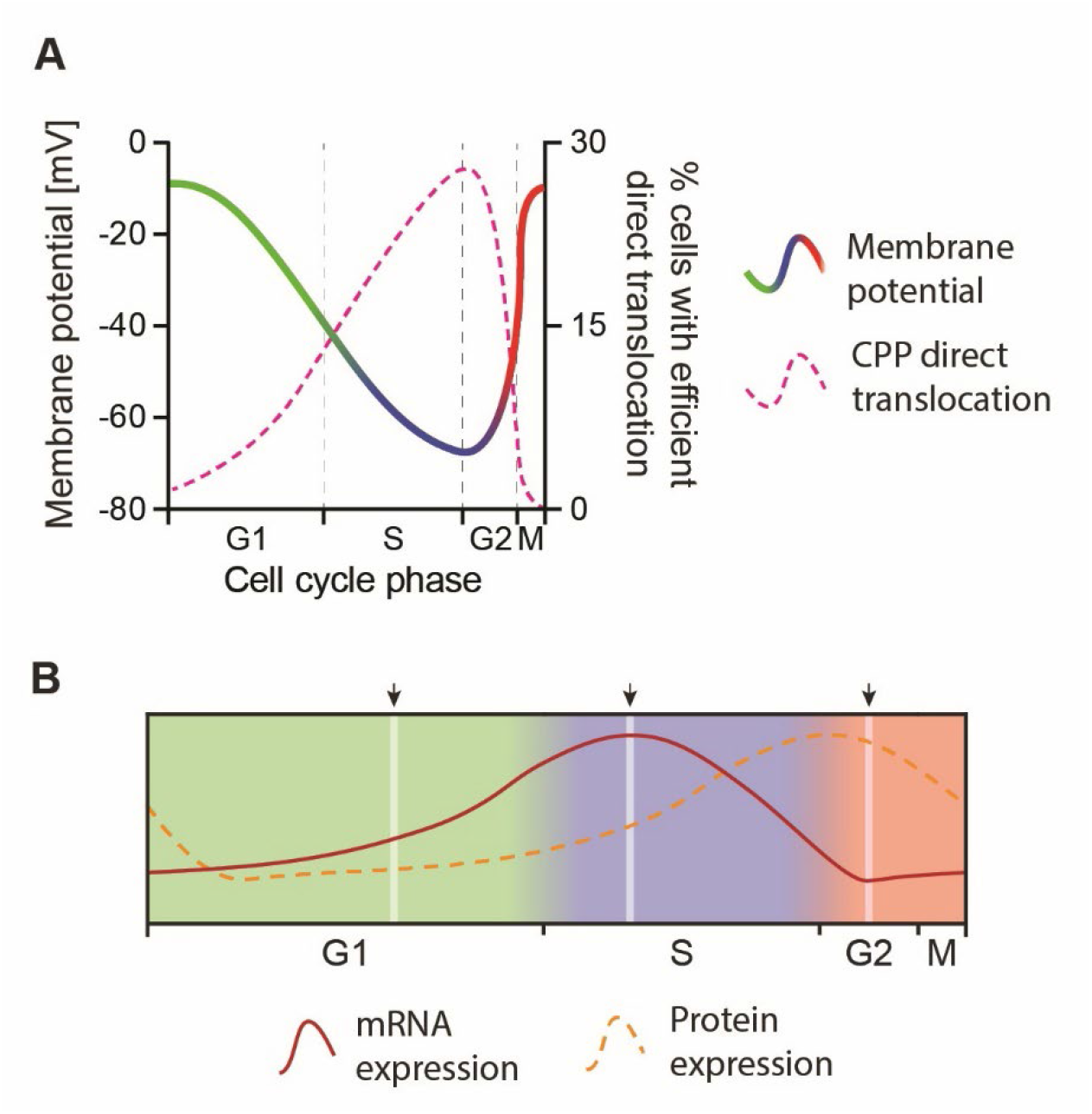
KCNH1 expression modulates the plasma membrane potential and CPP direct translocation. **(A)** The resting cell membrane potential oscillates throughout the cell cycle stages. G1 (approximate range: -5 to -45 mV) is characterized by a progressively hyperpolarizing cell membrane potential, which continues through the S phase (approximate range: -25 to -65 mV). Upon G2 onset (approximate range: -65 to -30 mV), membrane potential quickly depolarizes to values around -30 mV with the transition to early M phase. These variations of the cell membrane potential are correlated with the changes in CPP direct translocation, which is most efficient when cells are most hyperpolarized. **(B)** Schematic representation of measured KCNH1 mRNA levels (based on qPCR data from Figure 3C) and estimated KCNH1 protein levels across the cell cycle (shifted by 4 hours; see discussion). Arrows indicate collection points for KCNH1 mRNA.

KCNH1 expression is known to oscillate throughout the cell cycle and has been implicated in promoting cancer cell proliferation, apparently through its ability to regulate membrane potential during cell cycle progression [1, 31, 34]. Taking into account the estimated delay of at least two hours between protein transcription and the production, trafficking, and membrane insertion of functional channels [35], we propose that plasma membrane KCNH1 abundance is highest during the S-to-G2 transition and declines to a minimum during the G2-to-M transition (*Figure 4B*). Based on this model, increased KCNH1 activity during S/G2 promotes K⁺ efflux, contributing to progressive membrane hyperpolarization (*Figure 3A*), whereas the subsequent reduction in channel abundance during G2/M facilitates membrane depolarization (*Figure 3A*). Because membrane potential has emerged as an important determinant of CPP translocation[2–6, 18–22, 28], these cell cycle-dependent changes in KCNH1 expression provide a likely mechanistic link between cell cycle progression and the enhanced direct CPP uptake observed in synchronized cells, which peaks during G2 (*Figure 2*).

Our findings suggest that the cell cycle serves as a predictive marker for CPP uptake efficiency. Differential CPP cytosolic uptake along the cell cycle has important implications for the development of CPP clinical applications. For example, when targeting rapidly dividing cells such as cancer cells, a detailed understanding of their cell-cycle dynamics and membrane potential can guide the rational design of CPP delivery methods to improve efficacy and selectivity. Cancer cells are known to be globally depolarized [36], which should not favour CPP uptake. However, since cancer cells continuously divide, they go through S and G2 phases, and consequently through transient hyperpolarization states, more frequently than other cells, making these rapidly dividing cells good targets for CPP-derived therapeutics. We believe that these findings should be taken into consideration when investigating CPPs as potential delivery vectors.

## Material and Methods

### Cell lines

All cell lines were cultured in 5% CO2 at 37°C. HEK293t (ATCC: CRL-3216), U2OS ( ATCC: HTB-96), PIP-FUCCI (a generous gift from Jeffrey Jones), LAN1 (ACC655; Leibniz Institute DSMZ-German Collection of Microorganisms and Cell Cultures) were cultured in DMEM (61965-026; Thermo Fisher) supplemented with 10% heat-inactivated fetal bovine serum (FBS; A5256701; Thermo Fisher). All cell lines were mycoplasma negative.

### Peptides

Cy5-coupled R9 (RRRRRRRRR) and Cy5-TAT (RRRQRRKKRG) were synthesized in the retro-inverso conformation by SB-PEPTIDE, Lyon, France.

### Confocal microscopy

Confocal microscopy experiments were done on live cells. Cells were seeded onto glass bottom culture dishes (P35G-1.5-14-C; MatTek, corporation) and treated as described in the figures. Image acquisition was performed with a HC PL APO 63x/1.40 oil CS2 objective mounted on an inverted Leica Stellaris 8 laser scanning fluorescence confocal microscope equipped with four lasers (405nm DMOD, 488nm, 561nm, 638nm). Image resolution was 0.12 μm x 0.12 μm per pixel and pinhole size was set to maximum 1 AU or below. Cell images were acquired at a focal plane near the middle of the cell cutting through the nucleus either based in nuclear protein expression of PIP-FUCCI cells or incubation in presence of Hoechst 33342 (CDS023389, Sigma) of wild type cells. Images were acquired with 8-bit resolution (zoom: 0,75x). For detector HyD 1 (mTurquoise2), the following parameters were used: excitation laser 405 nm, laser intensity 10%, gain 10, spectral range acquisition 430 nm – 510 nm. For detector HyD 2 (mVenus), the following parameters were used: excitation laser 488 nm, laser intensity 15%, gain 5, spectral range acquisition 510 nm – 540 nm. For detector HyD 3 (mCherry), the following parameters were used: excitation laser 561nm, laser intensity 15%, gain 5, spectral range acquisition 580 nm – 627 nm. For detector HyD 4 (Cy5), the following parameters were used: excitation laser 638 nm, laser intensity 2%, gain 2, spectral range acquisition 645 nm – 750 nm. Images were collected using Leica’s LasX software.

### KCNH1 knock down using siPOOLs

60’000 U2OS cells were plated in 1,5 ml of DMEM with 10%FBS. The first round of siRNA transfection was performed right after the plating when the cells were still in suspension. The first transfection mix was made as follows: two 1.7 mL Eppendorf tubes were prepared per treatment, the first one containing 246 μL OptiMEM (Thermo 11058021) and 4 µL Lipofectamine RNAiMAX transfection reagent (Invitrogen 13778150); and the second one containing 210 µL OptiMEM and 40 µL siPOOLs siRNA, either non-targeting (siNeg), or targeting KCNH1 (siKCNH1). The tube was gently mixed and incubated at room temperature for 15 min. This mixture was added dropwise in the well containing the cells. One day later, the media was replaced with fresh DMEM+10%FBS, and the second transfection was performed, using 2 µL Lipofectamine 2000 transfection reagent (Invitrogen 11668019) instead of 4 µL Lipofectamine RNAiMAX. The cells were analysed 72 h after the first transfection, after which they were washed and treated with CPP as previously described in the methods. The final siRNA concentration in the media was 3 nM. The pools of 30 siRNAs targeting KCNH1, and the siRNA negative control (a pool of 30 nontargeting siRNA sequences), were purchased from the siTOOLS Biotech company.

### Synchronization of cells in G1/S

U2OS PIP-FUCCI, used for the membrane potential experiments and U2OS wild-type cells, used for CPP uptake visualization were plated at a concentration of 120’000 cells per well and incubated overnight in 5%CO2 at 37°C. PIP-FUCCI cells were plated on glass coverslips in a 6-well plate and wild-type U2OS cells were plated on 35-mm glass-bottom dishes in DMEM supplemented with 10% FBS. Cells were incubated with 2 mM thymidine for 18 hours, then washed with PBS, media was replaced by fresh media and incubated for 9 hours. After this period, a second 2 mM thymidine incubation lasting 18 hours was performed. Cells were released by washing with PBS and replacing it with fresh media and allowed to progress. Membrane potential measurements using whole cell patch clamp were then performed on the cells at the desired time points.

### Synchronization of cells in G2/M

U2OS PIP-FUCCI, used for the membrane potential experiments and U2OS wild-type cells, used for CPP uptake visualization were plated at a concentration of 250’000 cells per well and incubated overnight in 5%CO2 at 37°C. PIP-FUCCI cells were plated on glass coverslips in a 6-well plate and wild-type U2OS cells were plated on 35-mm glass-bottom dishes in DMEM supplemented with 10% FBS. Cells were preincubated with nocodazole for 24 hours, then washed with PBS and media was replaced by media containing 4 μM nocodazole (M1404, SIGMA-ALDRICH) alone or nocodazole and 15 μM CPP. Cells were incubated for 5 minutes and then visualized under a confocal microscope or were maintained in the media containing nocodazole during the membrane potential measurements. Confocal microscopy was performed instead of flow cytometry to quantify direct translocation in both conditions since the combination of CPP incubation and trypsinization caused 80% cell death. Samples containing DMSO alone were used as a negative control.

### Quantitation of cytosolic fluorescence

CPP cytosolic uptake was quantified based on confocal microscopy images. For CPP cytosolic uptake analysis, (Cy5 channel), fluorescence intensity signal reported in the figures was calculated by subtracting the background signal (coming from the dish) from the cytosolic signal recorded within the cells. Values higher than 0 a.u. were considered positive for direct translocation. The region of interest was selected within the region of cell cytosol devoid of endosomes.

### Membrane potential depolarization

Cells membrane potential was depolarized as by incubating cells for 10 min with 4 µg/ml gramicidin or by incubating cells for 30 min with depolarizing media [5]. Cells were then incubated with 15 µM CPP for 5 minutes at 37°C, washed in PBS and resuspended in FACS buffer (0.5mM EDTA, 1% FBS, in PBS – pre-filtered using PES membrane 0.22 µm filter, COBETTER LAB, Ref: SFM33PE0022S) prior to flow cytometry analysis using FACSARIA II SORP apparatus.

### KCNH1 inhibition with astemizole

U2OS wild-type cells were plated at a concentration of 250’000 cells per well and incubated overnight in 5%CO2 at 37°C either on glass coverslips in a 6-well plate and wild-type U2OS cells were plated on 35-mm glass-bottom dishes in DMEM supplemented with 10% FBS. Cells were then washed with PBS and media was replaced by media containing 10 µM astemizole (A2861, SIGMA-ALDRICH) alone or together with 15 µM CPP. Cells were incubated for 5 minutes and then visualized under a confocal microscope or were maintained in the media containing nocodazole during the membrane potential measurements. Samples containing DMSO alone were used as a negative control.

### Analysis of the phases of cell cycle (Hoechst and FUCCI)

#### PIP-FUCCI cells confocal microscopy

Image analysis was performed using Fiji software (version ImageJ 1.53t). PCNA analysis (mTurquoise2 channel) was performed visually, according to whether puncta were present in the nuclei. For PIP and Geminin1-110 analysis (mVenus and mCherry channels, respectively), intensity was measured in the nuclei. In each cell, across all channels (mTurquoise2, mVenus, mCherry) the intensities of the same 3 ROIs (background, corresponding to a region outside the cell; cytosol, corresponding to a region in the cytosol not containing cycle stage was then attributed to each cell depending on presence or absence of PCNA puncta, and on the signal intensity of both PIP and Geminin.

#### PIP-FUCCI cells flow cytometry

300’000 PIP-FUCCI cells were plated in suspension plates and incubated with 15 µM fluorescently-labelled CPP for 5 minutes at 37°C, after which they were centrifuged, and the supernatant removed, followed by PBS 1x wash. The cells were then pelleted once again, followed by resuspension in FACS buffer (0.5mM EDTA, 5% FBS, in PBS – pre-filtered using PES membrane 0.22 µm filter, COBETTER LAB, Ref: SFM33PE0022S). Single viable cells were then gated according to the PIP-FUCCI reporters (previously determined in the absence of CPPs) and CPP internalization in each group/cell stage (G1, S or G2/M) was assessed. Cells with efficient CPP direct translocation were arbitrarily defined, before gating the cells into given cell cycle stages, as the 10% brightest cells for the CPP signal (median: 76500 a.u., standard deviation: 27518.2 a.u.). The CPP fluorescence value defining those 10% brightest cells is reported on the figures as a dotted line.

#### DNA content based on Hoechst staining

80’000 U2OS wild-type cells were plated in 6-well plates and synchronized following the double thymidine block procedure described previously (see Synchronization of cells in G1/S). After this, cells were trypsinized and incubated with 2 µg/mL Hoechst-33342 for 30 minutes at 37°C. Flow cytometry was then performed using a CytoFLEX S (Beckman Coulter) apparatus, and 50’000 cells were recorded per time point.

### Cell sorting according to the CPP uptake efficiency

300’000 U2OS wild-type cells were plated in suspension plates and incubated with 15 μM fluorescently-labelled CPP for 5 minutes at 37°C, after which cells were centrifuged, and the supernatant removed, followed by PBS 1x wash. The cells were then pelleted once again, followed by resuspension in FACS buffer (0.5mM EDTA, 5% FBS, in PBS – pre-filtered using PES membrane 0.22 µm filter, COBETTER LAB, Ref: SFM33PE0022S). Profiling of the cell population was used to select three subpopulations with distinct direct translocation efficiencies (see *Figure 1 – Figure Supplement 1A*). Cells in these subpopulations were sorted into corresponding tubes containing RNALater transport/storage solution (RNAlater®, SIGMA-ALDRICH, MDL: MFCD03453003, Ref: R0901) for stabilization and protection of the RNA contents of the cells. This experiment included 4 replicates. RNA isolation was performed using Quiagen kit according to manufacturer’s instructions. Samples were sent for RNA sequencing, which was carried out by Illumina RNA-Seq.

### Transcriptomics data analysis

#### Data processing

Purity-filtered reads were adapters and quality trimmed with Cutadapt (v. 2.5, Martin 2011). Reads matching to ribosomal RNA sequences were removed with fastq_screen (v. 0.11.1). Remaining reads were further filtered for low complexity with reaper (v. 15-065, Davis et al. 2013). Reads were aligned against *Homo sapiens* GRCh38.102 genome using STAR (v. 2.5.3a, Dobin et al. 2013). The number of read counts per gene locus was summarized with htseq-count (v. 0.9.1, Anders et al. 2014) using gene annotation. Quality of the RNA-seq data alignment was assessed using RSeQC (v. 2.3.7, Wang et al. 2012). The htseq-generated counts data was used for the analysis.

#### Normalization and data transformation

Statistical analysis was performed in R (R version 4.2.2). Genes with low counts were filtered out according to the rule of 1 count per million (cpm) in at least 1 sample. Library sizes were scaled using TMM normalization. Subsequently, the normalized counts were transformed to cpm values and a log2 transformation was applied by means of the function cpm with the parameter setting prior.counts = 1 (EdgeR v 3.30.3; Robinson et al. 2010).

#### PCA analysis

After data normalization, a quality control analysis was performed through samples correlation/clustering and Principal Components Analysis (PCA). Batch-effect correction was applied using removeBatchEffect from the R package limma (Ritchie et al. 2015).

#### Statistical analysis with limma

DE analysis was performed on the batch-corrected log2 of the normalized CPM scores. The number of genes kept in the set after low counts filtering, and used for further analysis, is 16518. Differential expression was computed with the R Bioconductor package limma by fitting data to a linear model. P-values were adjusted using the Benjamini-Hochberg (BH) method, which controls for the false discovery rate (FDR).

After testing the DE genes for a FDR < 0.05, the genes were also filtered according to their fold-change. The fold-change cutoff was set to 1.5.

### Membrane potential measurements

250’000 U2OS PIP-FUCCI cells were plated overnight on glass coverslips in 6-well plates in DMEM +10%FBS and incubated in 5% CO2 at 37°C. The cells were then treated as indicated in the figures. The coverslips were transferred to bath solution. The cells were maintained at RT and were perfused with oxygenated bath solution at 9.9 ml/min. The bath solution contained (in mM): 130.9 NaCl, 5.2 KCl, 25.9 NaHCO3, 2 CaCl2, 1.2 MgCl2, 5.2 KCl, 1.25 NaH2PO4, 18 glucose and 1.7 ascorbic acid. The pipette was top filled with intracellular solution (295 mOsm) containing (in mM): 140 KGlu, 10 KCl, 10 HEPES, 10.0 phosphocreatine, 4.0 ATP-Mg2+, 0.3 GTP-Tris, 0.1 EGTA, with adjusted pH to 7.3. Whole-cell patch configuration was achieved by applying negative pressure on the patch pipette until seal resistance of at least 1 GΩ were obtained, after which pipette capacitance was compensated. Constant gentle negative pressure (suction) was then applied to gain cell access. The membrane potential recordings were performed in current clamp at 0 pA. The values recorded from cells with stable membrane potential signal over two minutes are displayed in the figures. The recordings were performed with 10 kHz frequency using the Multiclamp 700B Commander (Axon) and data was recorded using Clampex 11.2. Signal amplification was achieved with Digidata 1550B (Axon) amplifier. Analysis was performed using Clampfit 11.2. The patch setup was also equipped with an iXon X3 camera (Andor) along with SOLIS 64-bit (Andor) camera acquisition software, which allowed for image visualization, along with a blue laser (470 nm) and the U-MWBV2 filter cube. This also allowed us to distinguish cells in S phase based on punctate signal in the nucleus, thereby allowing us to have a control for the enrichment of S vs non-S cells in the synchronized population.

### RT-qPCR

Following the treatment indicated in the figures, cells were trypsinized and resuspended in RNALater transport/storage solution (RNAlater®, SIGMA-ALDRICH, MDL: MFCD03453003, Ref: R0901), and frozen at -20°C until processing. Total RNA was extracted from cells using RNeasy Mini Kit (Ref: 74106, QIAGEN) according to the manufacturer’s instructions. Reverse transcription from RNA to cDNA was performed using the RevertAid First Strand cDNA Synthesis Kit (Ref: K1622, THERMO SCIENTIFIC). Semiquantitative real-time PCR was performed using SYBR™ Select Master Mix for CFX (Ref: 4472942, APPLIED BIOSYSTEMS) using gene-specific primers (GAPDH-Forward: GAAGGTGAAGGTCGGAGT; GAPDH-Reverse: GAAGATGGTGATGGGATTTC; KCNH1-Forward: TAGTGGCCCCTCAAAACACG; KCNH1-Reverse: TTCTGCCCTGTGATAGCCAG). Data were normalized to mRNA levels of *GAPDH*, a housekeeping gene, and were analysed by the 2^−ΔΔCt^ method.

### Quantitation and Statistical Analysis

Statistical analysis was performed on non-normalized data, using GraphPad Prism 10.2. All measurements were from biological replicates. The details of statistical tests applied for the analysis and the exact number of cells quantified per condition can be found in figure legends.

## Acknowledgements

The laboratory of CW is supported by the Swiss National Science Foundation (grant n° 310030_207464) and AL by the SNSF grant (310030_214851) and Etat de Vaud. We are thankful to the Cellular Imaging Facility, the Genomic Technologies Facility and the Flow Cytometry Facility at the University of Lausanne for the resources provided and their technical help. We are grateful to Dr. Luis Pardo for his dedicated and persistent efforts to detect KCNH1 protein levels.

## Author contribution

Conception and design of study: FR, ET and CW

Acquisition of data: FR, ET, AH, and RM

Funding acquisition: AL and CW

Resources: AL and CW

Analysis and/or interpretation of data: FR, ET, AL and CW

Drafting the manuscript: FR, ET and CW

Revising the manuscript and approval of the submitted version: all authors.

## Data Availability

The transcriptomics datasets generated during the current study are available in the NCBI repository: https://www.ncbi.nlm.nih.gov/sra/?term=PRJNA1191216.

## Additional files

**Figure 1 – Figure Supplement 1.**
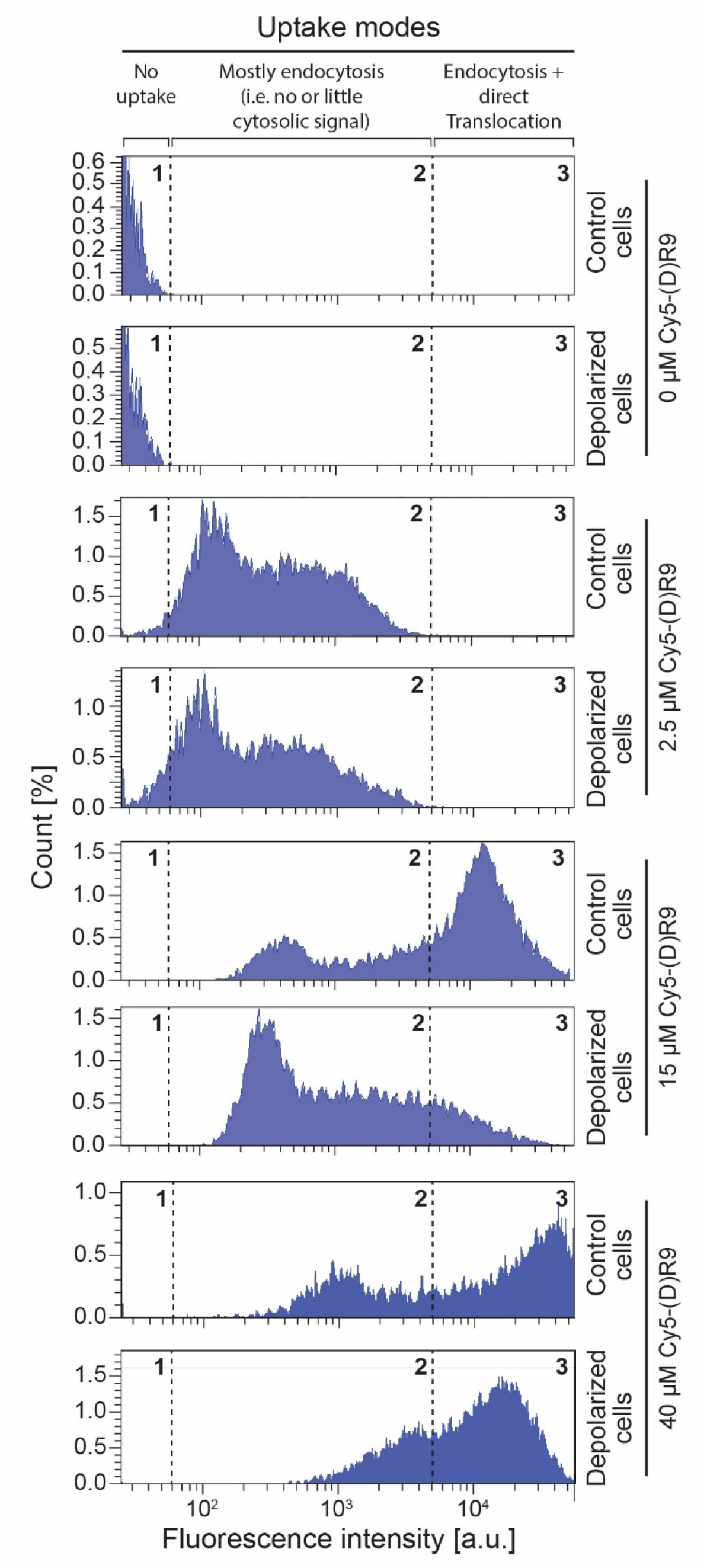
CPP modes of entry. Flow cytometry of wild-type U2OS cells incubated for 5 minutes at 37°C with different concentrations of Cy5-R9 in either control or high potassium concentration medium (depolarization medium). At low CPP concentrations (i.e. 2.5 μM), CPP-associated signal is found in intensity range 2 and is not affected by cell depolarization. Intensity range 2 represents CPP acquisition by endocytosis [37]. At 15 μM, the CPP cellular signal is also found in intensity range 3. This signal is strongly decreased by cell depolarization, indicating that most of the signal in intensity range 3 corresponds to CPP direct translocation [5]. The remaining signal in intensity range 3 after cell depolarization can be accounted for by the combined signal from endosomes and residual direct translocation. At very high CPP concentrations (i.e. 40 μM), the high potassium concentration medium only leads to a partial decrease in cell-associated CPP signal. At such high CPP concentrations, even when the potassium ion gradient is disrupted, the accumulation of the CPP-carried positive charges at the cell membrane is likely high enough to maintain sufficient polarization of the plasma membrane allowing some direct translocation to occur [5].

**Figure 1 – Figure Supplement 2.**
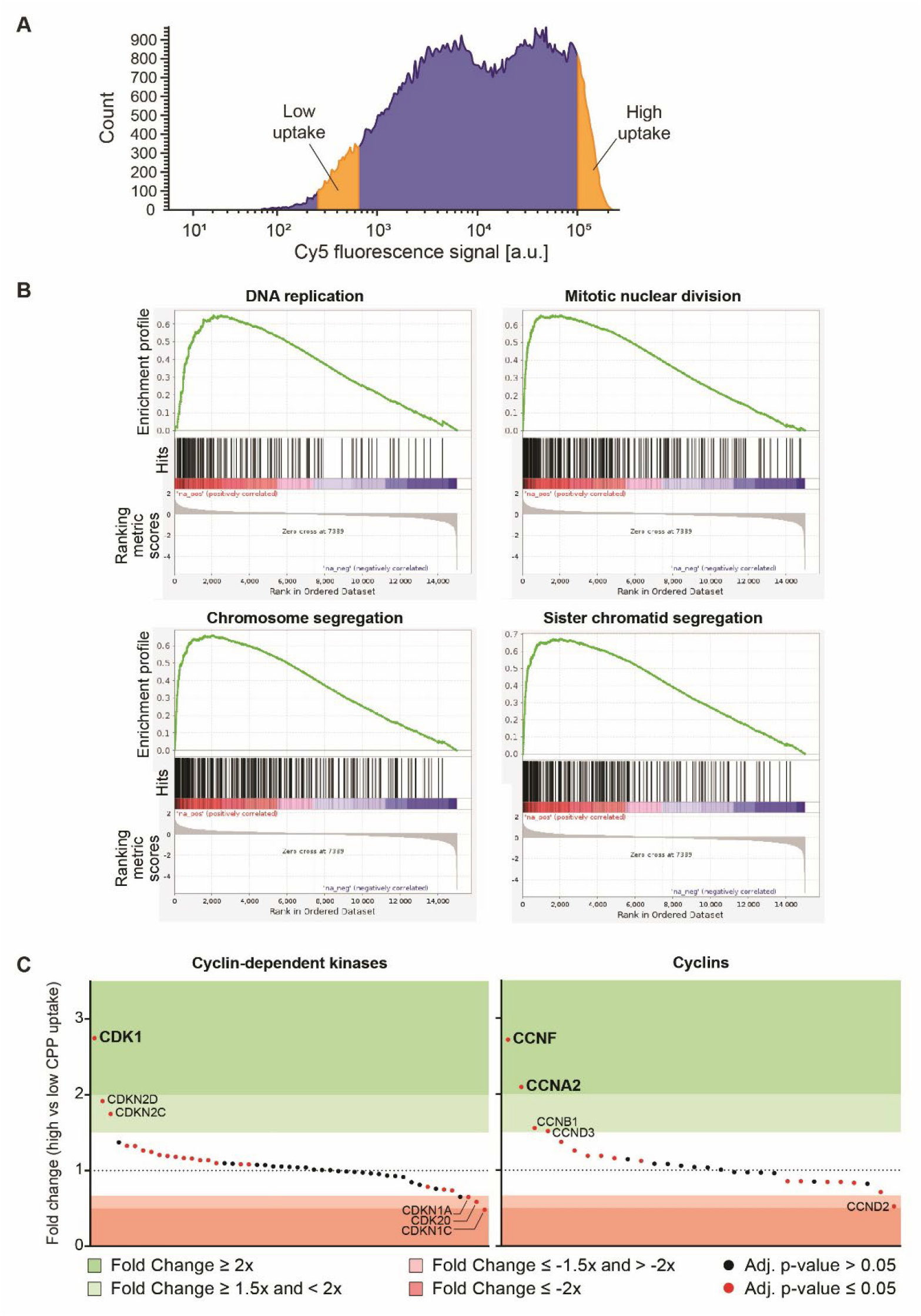
Differential gene expression between high- and low- CPP cytosolic uptakers. **(A)** Wild-type U2OS cells were incubated with 15 µM Cy5-R9 for 5 minutes at 37°C and then washed. The 5% dimmest and 5% brightest cells (indicated in yellow in the figure) were sorted flow cytometry. From these two subpopulations, RNA was isolated and used for transcriptomics analysis. **(B)** Gene set enrichment analysis of the four top gene sets enriched in cells with high CPP uptake. **(C)** Differential expression of cyclins and cyclin-dependent kinases in cells with high vs low CPP cytosolic uptake. Cyclin-dependent kinase family members (left panel) and cyclin family members (right panel) were ranked along the x axis according to the fold change of their mRNA expression in cells with high vs low CPP uptake.

**Figure 2 – Figure Supplement 1.**
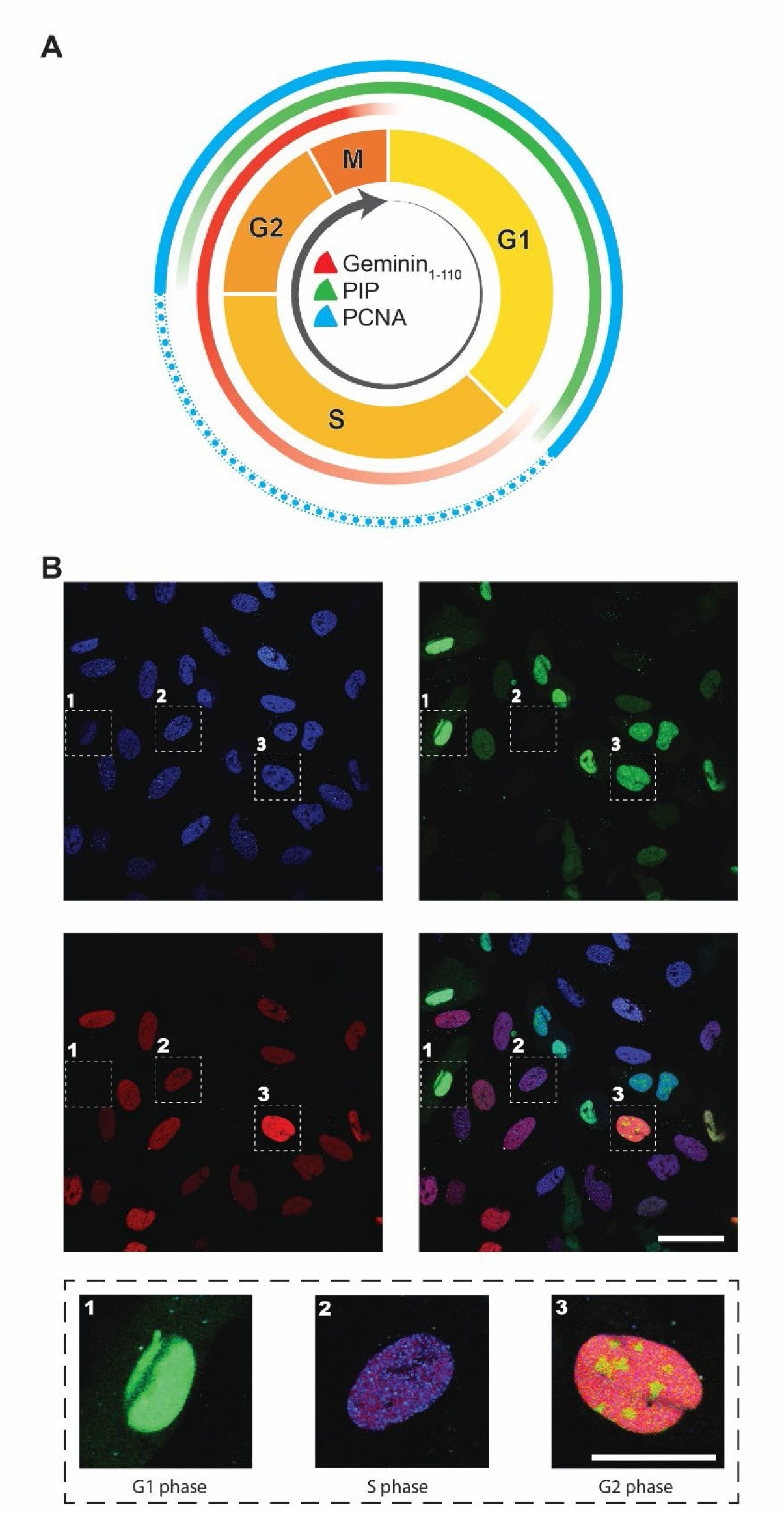
The PIP-FUCCI reporter system. **(A)** U2OS PIP-FUCCI cells express three differentially colored markers that label the cells according to the cell cycle phase they are in. The expression of these markers along the cell cycle is schematically indicated in the figure. The rationale of using these three markers to follow progression of cells throughout the cell cycle is explained below. **mCherry-Geminin1-110** (represented in red). After DNA replication is initiated, it is critical that origins of replications do not refire. Geminin is one protein that ensures this. Geminin accumulates in the nucleus in early S phase, binding to and sequestering Cdt1. This prevents Cdt1 from binding to the minichromosome maintenance (MCM) complex and to DNA (the MCM complex is part of the pre-replication complex). Consequently, the MCM complex cannot be reloaded onto the replicated chromatin and re-replication is inhibited. From the end of mitosis and throughout G1, Geminin is degraded by the APC/C (anaphase-promoting complex/cyclosome) complex, permitting pre-replication complex assembly for the next cycle of DNA replication. It is the N-terminal part of Geminin that is targeted by the APC/C complex (sequence 1-110). **PIP-mVenus** (or **Cdt11-17-mVenus**) (represented in green). The Cdt1 protein is required for pre-replication complex activity. To prevent DNA re-replication, cells must make sure that Cdt1 does not operate during S phase. This can be achieved through Geminin-mediated inhibition (see above) but also through degradation by the proteasome. Proteasome-mediated Cdt1 degradation can proceed in two distinct ways: i) following its phosphorylation by the cyclin A/CDK1 and cyclin A/CDK2 complexes, Cdt1 is ubiquitinated by the SCF^Skp2^ ubiquitin ligase and degraded; ii) upon binding to PCNA (proliferating cell nuclear antigen) on chromatin, Cdt1 is degraded by the Cul4-DDB1^Cdt2^ ubiquitin ligase. The Cdt1 part that is recognized by PCNA is called PIP (PCNA-interacting protein) and corresponds to the first 17 amino acids of the protein. **mTurquoise2-PCNA** (represented in blue). When PCNA binds to chromatin, it interacts with Cdt1 that is then degraded. PCNA binding to chromatin forms puncta. This occurs during S phase. Outside of S phase, PCNA is not binding to DNA, is not interacting with Cdt1, and has a diffuse location in the nucleus. **(B)** Images of an asynchronous culture of U2OS PIP-FUCCI cells were taken using channels allowing the three markers to be specifically detected (Figure 2 – Supplement 2). The combination of colors allows for cell cycle phase determination: cells in G1 are labelled in green only (PIP-mVenus), cells in S phase are predominantly labelled in red (mCherry-Geminin1-110), and cells in G2/M are characterized by the presence of both signals. The third marker (mTurquoise2-PCNA), when seen as dots in the nucleus, is an accurate marker for S phase. Scale bar: 25 µm.

**Figure 2 – Figure Supplement 2.**
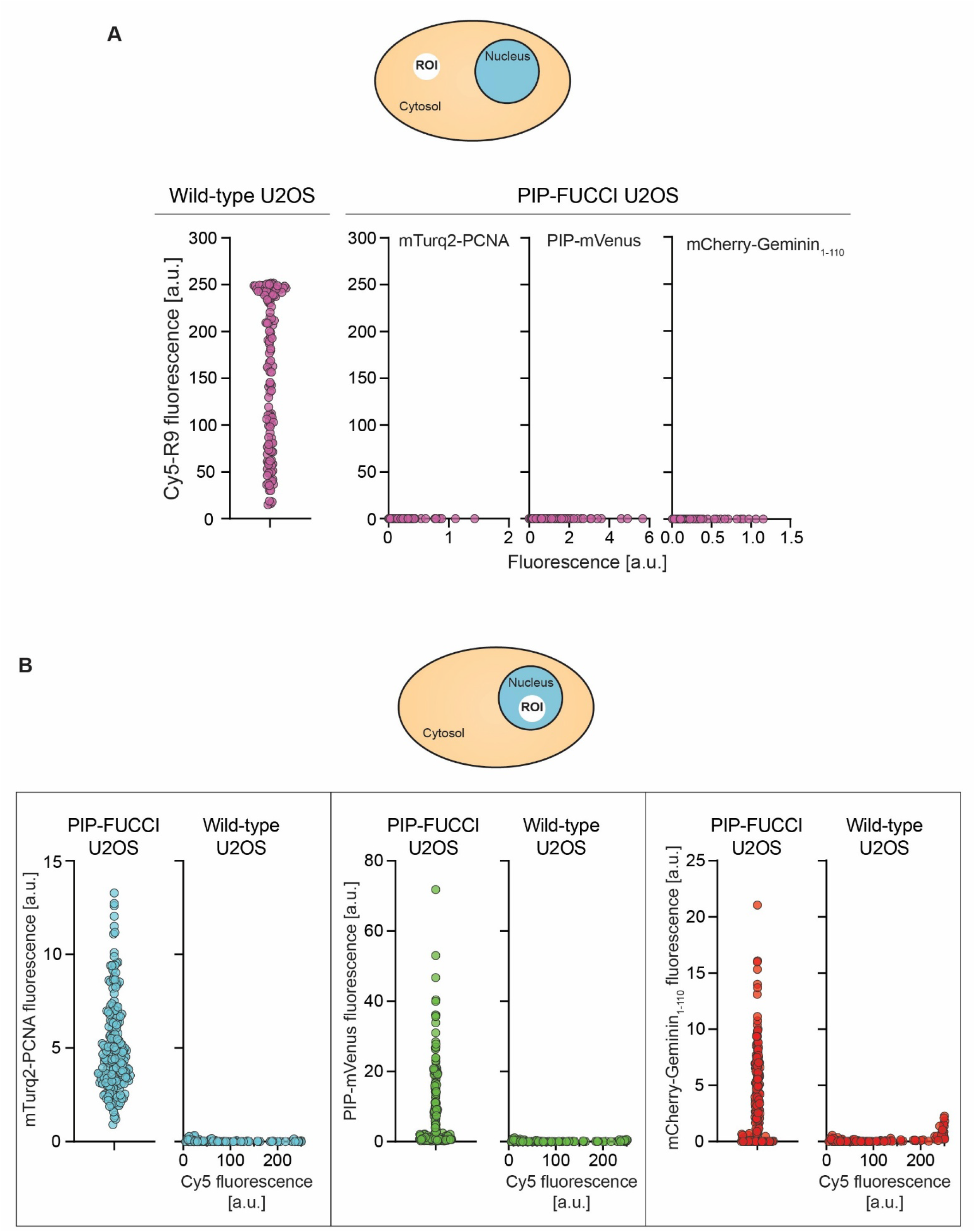
PIP-FUCCI markers and Cy5-R9 bleed-through assessment by confocal microscopy. **(A)** No bleed-through was detected of the Cy5 signal, carried by CPPs, into the PIP-FUCCI-specific marker channels. Left graph: Cy5-R9 cytosolic fluorescence in wild-type U2OS cells incubated with Cy5-R9 (n=126). Right three graphs: mTurq2-PCNA, PIP-mVenus and mCherry-Geminin1-110 fluorescence (*x axis*) and R9-Cy5 fluorescence (*y-axis*) from given cytosolic region-of-interest (ROI) (see cartoon on top) in PIP-FUCCI U2OS cells (n=196). **(B)** No bleed-through was detected of the PIP-FUCCI-specific marker signal into the Cy5 channel. In each panel, the graph on the left depicts the fluorescence of a given PIP-FUCCI marker in the nucleus (n = 196). In each panel, the graph on the right shows how much of the fluorescence of R9-Cy5 (x-axis) bleeds into a given PIP-FUCCI marker channel (y-axis) from given nuclear region-of-interest (ROI) (see cartoon on top) in PIP-FUCCI U2OS cells (n=126).

**Figure 2 – Figure Supplement 3.**
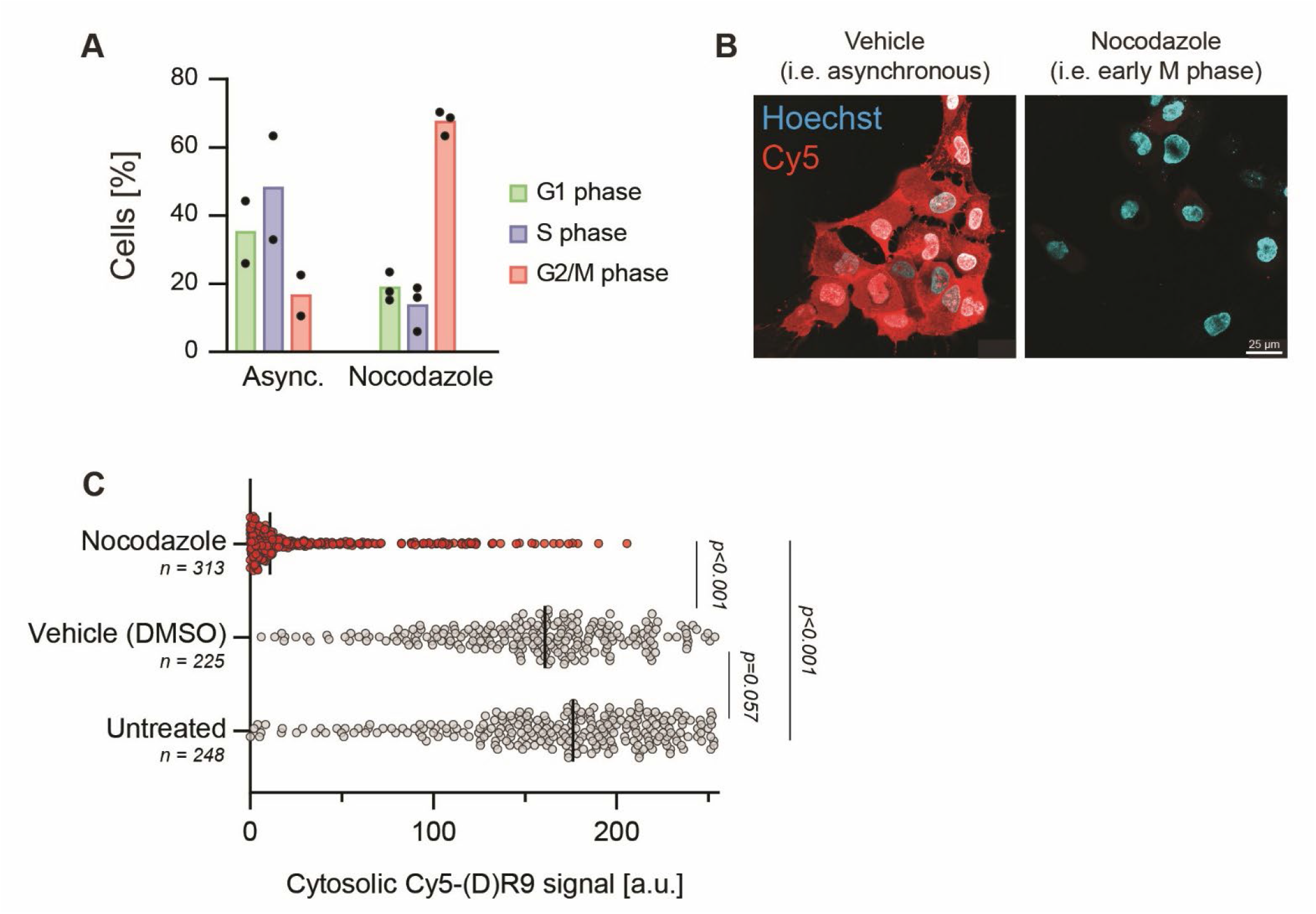
Mitotic cells do not uptake CPPs efficiently. **(A)** Nocodazole block leads to enrichment of cells in M phase. Quantitation of percentage of cells in specific cell cycle stages upon treatment with 4 μM nocodazole or vehicle (0.3% DMSO) for 24 hours at 37°C. Cell cycle stage was determined based on the expression of U2OS PIP-FUCCI-associated markers. **(B)** Representative confocal images of wild-type U2OS cells incubated with 15 µM Cy5-R9 for 5 minutes in cells preincubated either with 4 µM nocodazole or with a vehicle control (0.3% DMSO) for 24 hours at 37°C. Cells were then washed in PBS and fresh media was added. **(C)** CPP direct translocation is drastically blocked when cells are synchronized in early mitosis. U2OS cells were left untreated or incubated with 15 µM Cy5-R9 for 5 minutes in cells preincubated either with 4 µM nocodazole or with a vehicle control (0.3% DMSO) for 24 hours at 37°C. CPP cytosolic signal was quantitated based on confocal microscopy images from three independent experiments. Statistical significance is indicated and was calculated using Brown-Forsythe and Welch ANOVA tests.

**Figure 2 – Figure Supplement 4.**
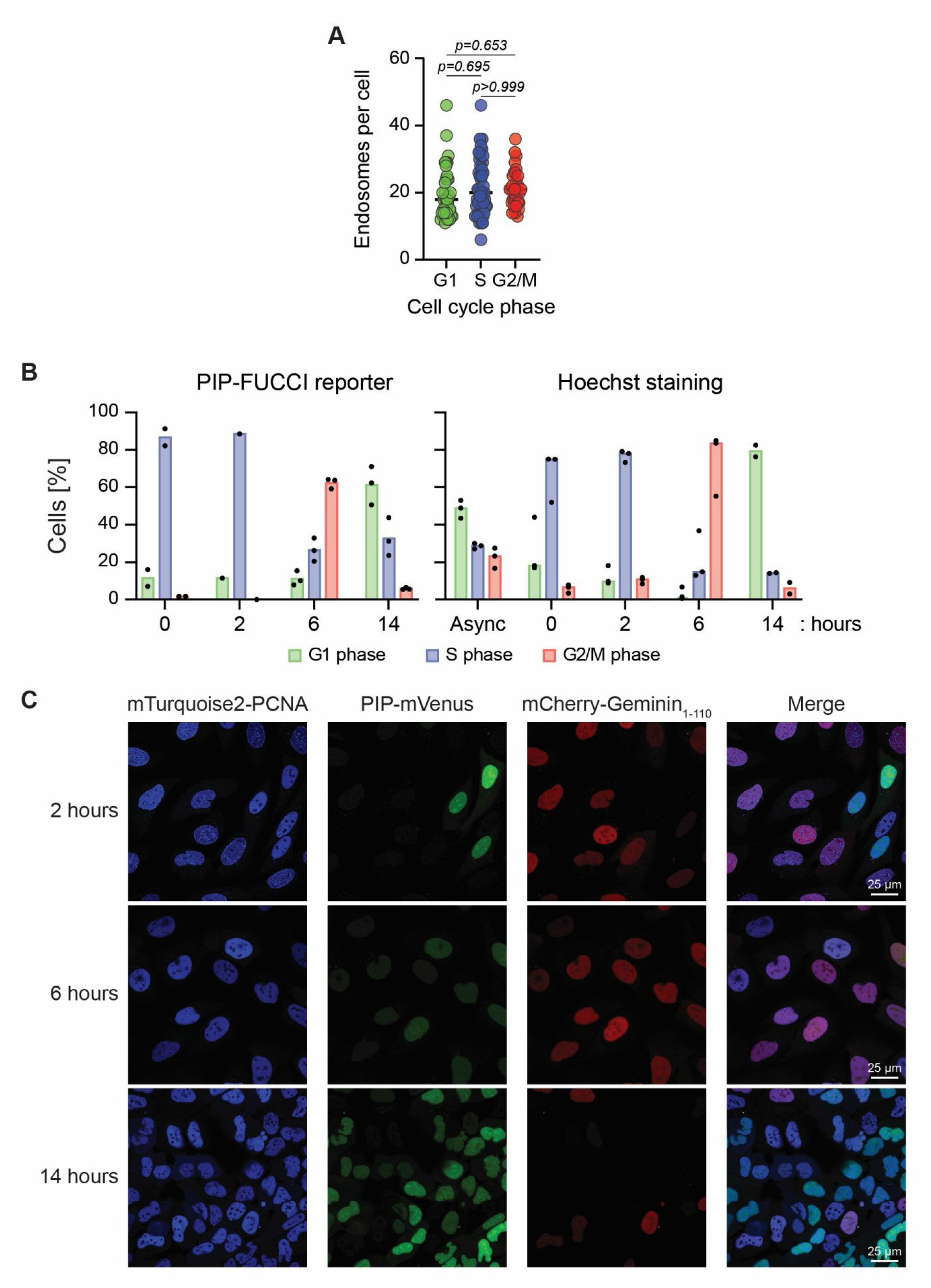
Cell cycle synchronization. **(A)** Endocytosis remains constant across the cell cycle. Endosome number quantitation per cell cycle stage based on confocal imaging of U2OS PIP-FUCCI cells incubated with R9-Cy5 for 5 minutes at 37°C. Statistical significance was assessed using Brown-Forsythe and Welch ANOVA tests (p-value G1vsS= 0.6954; p-value SvsG2/M>0.9999; p-value G1vsG2/M= 0.6525). **(B)** Double thymidine block allows for efficient synchronization of cells in S-phase and, upon release, to progression through specific stages of the cell cycle. Quantitation of percentage of cells at a given cell cycle stage at the indicated time points based on U2OS PIP-FUCCI-associated protein expression (left) or based on DNA content measured with Hoechst (right). **(C)** Representative confocal images of U2OS PIP-FUCCI cells synchronized by double thymidine block.

**Figure 3 – Figure Supplement 1.**
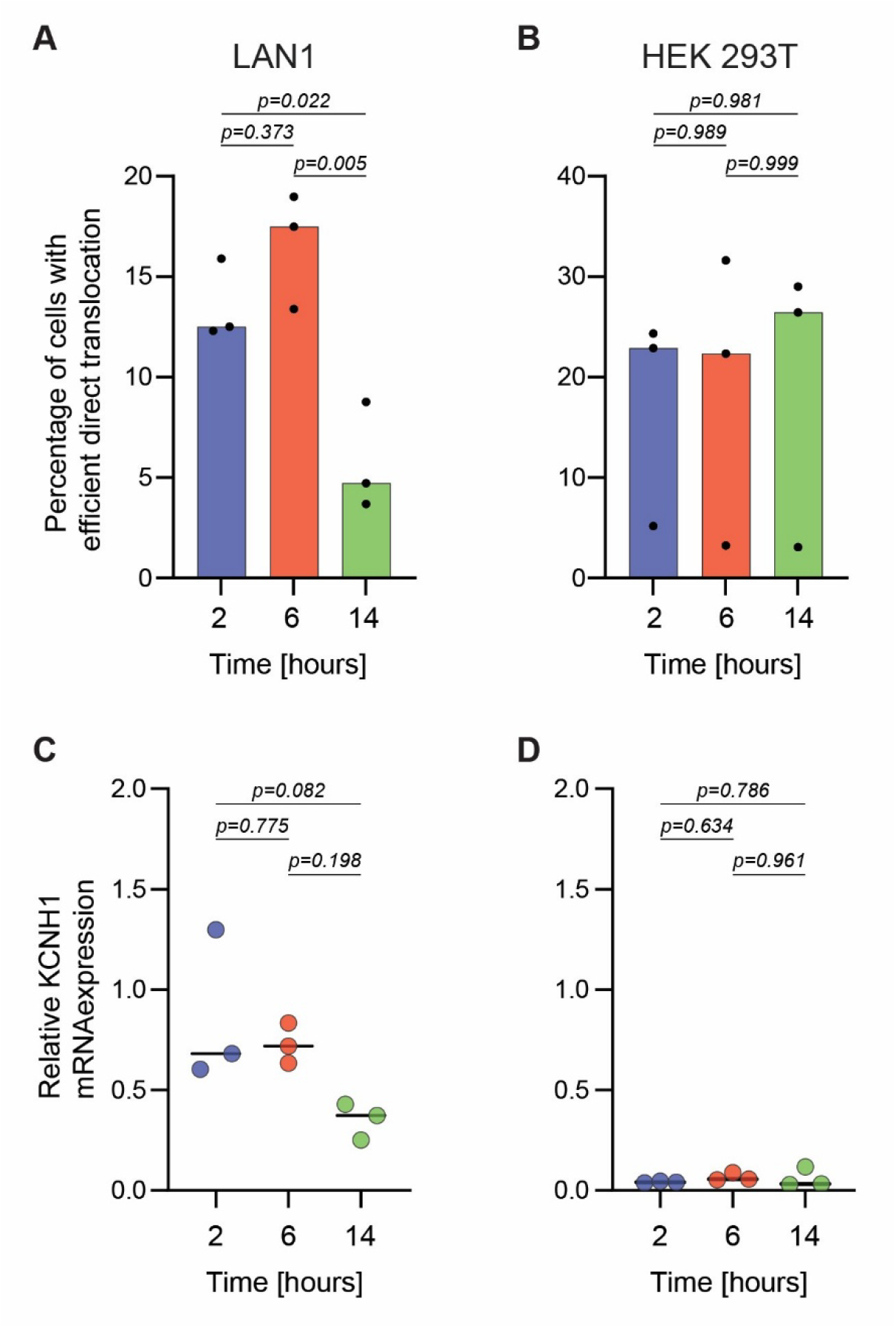
LAN1, but not HEK293T, cells exhibit cell cycle-dependent CPP cytosolic uptake. **(A)** Percentage of cells within a given cell cycle stage that acquired Cy5-R9 efficiently through direct translocation in LAN1 and HEK 293T cells synchronized using a double-thymidine block. **(B)** Semi-quantitative PCR assessment of KCNH1 mRNA levels in double thymidine-synchronized LAN1 and HEK 293T cells at different time points post release. Samples were normalized to GAPDH mRNA levels. Statistical analysis was performed using ANOVA multiple comparison with Tukey’s correction.

**Figure 3 – Figure Supplement 2.**
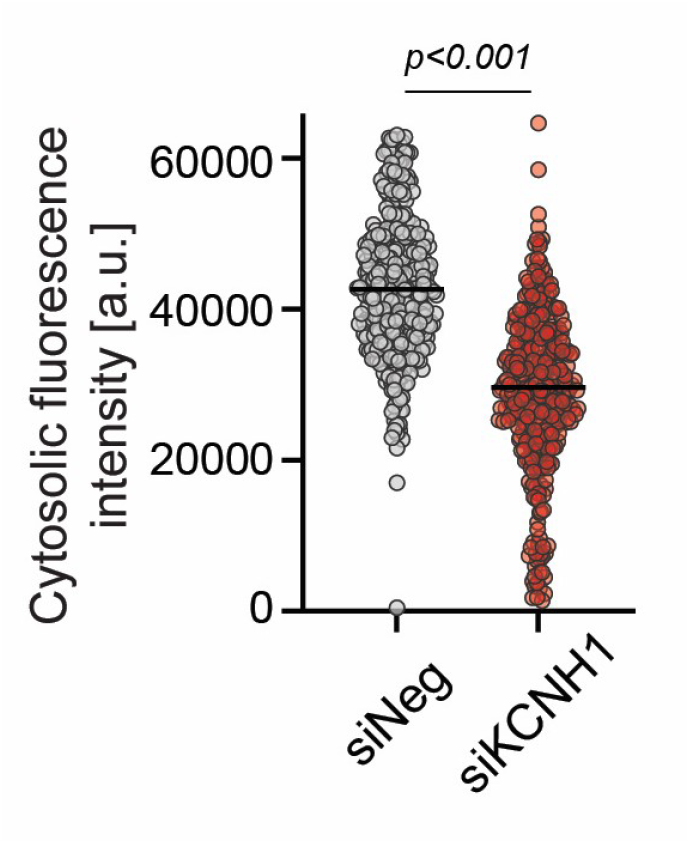
TAT-Cy5 internalization in U2OS cells is KCNH1-dependent. KCNH1 knock-down reduces TAT-Cy5 direct translocation. A total of 347 cells were analysed in three independent experiments. Statistical significance was performed using Welch’s t test.

**Table S1.** Depolarizing media composition.

| CAS Number | Component<br>s | Quantity in g/l |
| --- | --- | --- |
| 13477-34-4 | Calcium Nitrate Tetrahydrate | 0.10000000 |
| 7487-88-9 | Magnesium Sulfate Anhydrous | 0.04884000 |
| 50-99-7 | D-Glucose Anhydrous | 2.00000000 |
| 56-40-6 | Glycine | 0.01000000 |
| 39537-23-0 | L-Alanyl-L-Glutamine | 0.44600000 |
| 74-79-3 | L-Arginine Free Base | 0.20000000 |
| 70-47-3 | L-Asparagine Anhydrous | 0.05000000 |
| 56-84-8 | L-Aspartic acid | 0.02000000 |
| 30925-07-6 | L-Cystine Dihydrochloride | 0.06520000 |
| 56-86-0 | L-Glutamic Acid | 0.02000000 |
| 71-00-1 | L-Histidine | 0.01500000 |
| 51-35-4 | L-Hydroxy-L-Proline | 0.02000000 |
| 73-32-5 | L-Isoleucine | 0.05000000 |
| 61-90-5 | L-Leucine | 0.05000000 |
| 657-27-2 | L-Lysine Monohydrochloride | 0.04000000 |
| 63-68-3 | L-Methionine | 0.01500000 |
| 63-91-2 | L-Phenylalanine | 0.01500000 |
| 147-85-3 | L-Proline | 0.02000000 |
| 56-45-1 | L-Serine | 0.03000000 |
| 72-19-5 | L-Threonine | 0.02000000 |
| 73-22-3 | L-Tryptophan | 0.00500000 |
| 69847-45-6 | L-Tyrosine Disodium Salt Dihydrate | 0.02883000 |
| 72-18-4 | L-Valine | 0.02000000 |
| 67-48-1 | Choline Chloride | 0.00300000 |
| 58-85-5 | D-Biotin | 0.00020000 |
| 137-08-6 | D-Ca Pantothenate | 0.00025000 |
| 59-30-3 | Folic Acid | 0.00100000 |
| 87-89-8 | Myo-Inositol | 0.03500000 |
| 98-92-0 | Nicotinamide (Nicotinic acid amide) | 0.00100000 |
| 150-13-0 | P-Aminobenzoic Acid (PABA) | 0.00100000 |
| 58-56-0 | Pyridoxine Hydrochloride | 0.00100000 |
| 83-88-5 | Riboflavin | 0.00020000 |
| 67-03-8 | Thiamine Hydrochloride | 0.00100000 |
| 68-19-9 | Vitamin B12 | 0.00000500 |
| 70-18-8 | L-Glutathione Reduced | 0.00100000 |
| 34487-61-1 | Phenol Red Sodium Salt | 0.00530000 |
| WATER |  | 996.66117500 |

## References

1. Rao, V.R., M. Perez-Neut, S. Kaja, and S. Gentile, Voltage-gated ion channels in cancer cell proliferation. Cancers (Basel), 2015. 7(2): p. 849–75.

2. Rothbard, J.B., T.C. Jessop, R.S. Lewis, B.A. Murray, and P.A. Wender, Role of Membrane Potential and Hydrogen Bonding in the Mechanism of Translocation of Guanidinium-Rich Peptides into Cells. Journal of the American Chemical Society, 2004. 126(31): p. 9506–9507.

3. Wallbrecher, R., T. Ackels, R.A. Olea, M.J. Klein, L. Caillon, J. Schiller, P.H. Bovee-Geurts, T.H. van Kuppevelt, A.S. Ulrich, M. Spehr, M.J.W. Adjobo-Hermans, and R. Brock, Membrane permeation of arginine-rich cell-penetrating peptides independent of transmembrane potential as a function of lipid composition and membrane fluidity. J Control Release, 2017. 256: p. 68–78.

4. Zhang, X., Y. Jin, M.R. Plummer, S. Pooyan, S. Gunaseelan, and P.J. Sinko, Endocytosis and membrane potential are required for HeLa cell uptake of R.I.-CKTat9, a retro-inverso Tat cell penetrating peptide. Mol Pharm, 2009. 6(3): p. 836–48.

5. Trofimenko, E., G. Grasso, M. Heulot, N. Chevalier, M.A. Deriu, G. Dubuis, Y. Arribat, M. Serulla, S. Michel, G. Vantomme, F. Ory, L.C. Dam, J. Puyal, F. Amati, A. Luthi, A. Danani, and C. Widmann, Genetic, cellular, and structural characterization of the membrane potential-dependent cell-penetrating peptide translocation pore. Elife, 2021. 10.

6. Kuriyama, M., H. Hirose, Y. Kawaguchi, J. Michibata, M. Maekawa, and S. Futaki, KCNN4 as a genomic determinant of cytosolic delivery by the attenuated cationic lytic peptide L17E. Mol Ther, 2025. 33(2): p. 595–614.

7. Vasconcelos, L., K. Parn, and U. Langel, Therapeutic potential of cell penetrating peptides. Therapeutic delivery, 2013. 4(5): p. 573–91.

8. Madani, F., S. Lindberg, U. Langel, S. Futaki, and A. Graslund, Mechanisms of cellular uptake of cell-penetrating peptides. J Biophys, 2011. 2011: p. 414729.

9. Bechara, C. and S. Sagan, Cell-penetrating peptides: 20 years later, where do we stand? FEBS Lett, 2013. 587(12): p. 1693–702.

10. Wang, F., Y. Wang, X. Zhang, W. Zhang, S. Guo, and F. Jin, Recent progress of cell-penetrating peptides as new carriers for intracellular cargo delivery. J Control Release, 2014. 174: p. 126–36.

11. Frankel, A.D. and C.O. Pabo, Cellular uptake of the Tat protein from human immunodeficience virus. Cell, 1988. 55: p. 1189–1193.

12. Joliot, A., C. Pernelle, H. Deagostini-Bazil, and A. Prochiantz, Antennapedia homebox peptide regulates neural morphogenesis. Proceedings of the National Academy of Sciences of the United States of America, 1991. 88: p. 1864–1868.

13. Futaki, S., W. Ohashi, T. Suzuki, M. Niwa, S. Tanaka, K. Ueda, H. Harashima, and Y. Sugiura, Stearylated Arginine-Rich Peptides: A New Class of Transfection Systems. Bioconjugate Chemistry, 2001. 12(6): p. 1005–1011.

14. Delaroche, D., B. Aussedat, S. Aubry, G. Chassaing, F. Burlina, G. Clodic, G. Bolbach, S. Lavielle, and S. Sagan, Tracking a New Cell-Penetrating (W/R) Nonapeptide, through an Enzyme-Stable Mass Spectrometry Reporter Tag. Analytical Chemistry, 2007. 79(5): p. 1932–1938.

15. Guidotti, G., L. Brambilla, and D. Rossi, Cell-Penetrating Peptides: From Basic Research to Clinics. Trends Pharmacol Sci, 2017. 38(4): p. 406–424.

16. Xie, J., Y. Bi, H. Zhang, S. Dong, L. Teng, R.J. Lee, and Z. Yang, Cell-Penetrating Peptides in Diagnosis and Treatment of Human Diseases: From Preclinical Research to Clinical Application. Front Pharmacol, 2020. 11: p. 697.

17. Koo, J.-H., G.-R. Kim, K.-H. Nam, and J.-M. Choi, Unleashing cell-penetrating peptide applications for immunotherapy. Trends in Molecular Medicine, 2022. 28(6): p. 482–496.

18. Via, M.A., J. Klug, N. Wilke, L.S. Mayorga, and M.G. Del Popolo, The interfacial electrostatic potential modulates the insertion of cell-penetrating peptides into lipid bilayers. Phys Chem Chem Phys, 2018. 20(7): p. 5180–5189.

19. Lin, J. and A. Alexander-Katz, Cell membranes open “doors” for cationic nanoparticles/biomolecules: insights into uptake kinetics. ACS Nano, 2013. 7(12): p. 10799–808.

20. Gao, X., S. Hong, Z. Liu, T. Yue, J. Dobnikar, and X. Zhang, Membrane potential drives direct translocation of cell-penetrating peptides. Nanoscale, 2019. 11(4): p. 1949–1958.

21. Moghal, M.M.R., M.Z. Islam, F. Hossain, S.K. Saha, and M. Yamazaki, Role of Membrane Potential on Entry of Cell-Penetrating Peptide Transportan 10 into Single Vesicles. Biophys J, 2020. 118(1): p. 57–69.

22. Franke, J., P. Dubatouka, A. Yourdkhani, S. Soni, T. Utesch, J. Serrano, T. Soykan, M. Lehmann, H. Sun, J.V.V. Arafiles, and C.P.R. Hackenberger, Cellular protein delivery through membrane potential driven water pores. bioRxiv, 2026: p. 2026.02.24.707441.

23. Herce, H.D., A.E. Garcia, and M.C. Cardoso, Fundamental molecular mechanism for the cellular uptake of guanidinium-rich molecules. J Am Chem Soc, 2014. 136(50): p. 17459–67.

24. Mootha, V.K., C.M. Lindgren, K.F. Eriksson, A. Subramanian, S. Sihag, J. Lehar, P. Puigserver, E. Carlsson, M. Ridderstråle, E. Laurila, N. Houstis, M.J. Daly, N. Patterson, J.P. Mesirov, T.R. Golub, P. Tamayo, B. Spiegelman, E.S. Lander, J.N. Hirschhorn, D. Altshuler, and L.C. Groop, PGC-1α-responsive genes involved in oxidative phosphorylation are coordinately downregulated in human diabetes. Nature Genetics, 2003. 34(3): p. 267–273.

25. Subramaniana, A., P. Tamayoa, V.K. Moothaa, S. Mukherjeed, B.L. Eberta, M.A. Gillettea, A. Paulovichg, S.L. Pomeroyh, T.R. Goluba, E.S. Landera, and J.P. Mesirova, Gene set enrichment analysis: A knowledge-based approach for interpreting genome-wide expression profiles. PNAS, 2005. 102(43): p. 15545–15550.

26. Grant, G.D., K.M. Kedziora, J.C. Limas, J.G. Cook, and J.E. Purvis, Accurate delineation of cell cycle phase transitions in living cells with PIP-FUCCI. Cell Cycle, 2018. 17(21-22): p. 2496–2516.

27. Blajeski, A.L., V.A. Phan, T.J. Kottke, and S.H. Kaufmann, G1 and G2 cell-cycle arrest following microtubule depolymerization in human breast cancer cells. Journal of Clinical Investigation, 2002. 110(1): p. 91–99.

28. Trofimenko, E., N. Gervasi, S. Perez, N. Rodriguez, D. Ravault, S. Cribier, H. Berry, L. Venance, and S. Sagan, Transient pores account for cell-penetrating peptide and homeoprotein translocation. Proc Natl Acad Sci U S A, 2026. 123(26): p. e2602649123.

29. Serulla, M., P. Anees, A. Hallaj, E. Trofimenko, T. Kalia, Y. Krishnan, and C. Widmann, Plasma membrane depolarization reveals endosomal escape incapacity of cell-penetrating peptides. Eur J Pharm Biopharm, 2023. 184: p. 116–124.

30. Garcia-Ferreiro, R.E., D. Kerschensteiner, F. Major, F. Monje, W. Stuhmer, and L.A. Pardo, Mechanism of block of hEag1 K+ channels by imipramine and astemizole. J Gen Physiol, 2004. 124(4): p. 301–17.

31. Urrego, D., A. Sanchez, A.P. Tomczak, and L.A. Pardo, The electric fence to cell-cycle progression: Do local changes in membrane potential facilitate disassembly of the primary cilium?: Timely and localized expression of a potassium channel may set the conditions that allow retraction of the primary cilium. Bioessays, 2017. 39(6).

32. Bachmann, M., W. Li, M.J. Edwards, S.A. Ahmad, S. Patel, I. Szabo, and E. Gulbins, Voltage-Gated Potassium Channels as Regulators of Cell Death. Front Cell Dev Biol, 2020. 8: p. 611853.

33. Cone Jr., C.D., Electroosmotic interactions accompanying mitosis initiation in sarcoma cells in vitro. Section of Biological and Medical Sciences, 1968.

34. Urrego, D., A.P. Tomczak, F. Zahed, W. Stuhmer, and L.A. Pardo, Potassium channels in cell cycle and cell proliferation. Philos Trans R Soc Lond B Biol Sci, 2014. 369(1638): p. 20130094.

35. Petaja-Repo, U.E., M. Hogue, A. Laperriere, P. Walker, and M. Bouvier, Export from the endoplasmic reticulum represents the limiting step in the maturation and cell surface expression of the human delta opioid receptor. J Biol Chem, 2000. 275(18): p. 13727–36.

36. Capiod, T., Cell proliferation, calcium influx and calcium channels. Biochimie, 2011. 93(12): p. 2075–9.

37. Trofimenko, E., Y. Homma, M. Fukuda, and C. Widmann, The endocytic pathway taken by cationic substances requires Rab14 but not Rab5 and Rab7. Cell Rep, 2021. 37(5): p. 109945.

